# Trait-specific trade-offs prevent niche expansion in two parasites

**DOI:** 10.1101/621581

**Authors:** Eva J. P. Lievens, Yannis Michalakis, Thomas Lenormand

## Abstract

The evolution of host specialization has been studied intensively, yet it is still often difficult to determine why parasites do not evolve broader niches – in particular when the available hosts are closely related and ecologically similar. Here, we used an experimental evolution approach to study the evolution of host specialization, and its underlying traits, in two sympatric parasites: *Anostracospora rigaudi* and *Enterocytospora artemiae*, microsporidians infecting the brine shrimp *Artemia franciscana* and *Artemia parthenogenetica*. In the field, both parasites regularly infect both hosts, yet experimental work has revealed that they are each partially specialized. We serially passaged the parasites on one, the other, or an alternation of the two hosts; after ten passages, we assayed the infectivity, virulence, and spore production rate of the evolved lines. In accordance with previous studies, *A. rigaudi* maintained a higher fitness on *A. parthenogenetica*, and *E. artemiae* on *A. franciscana*, in all treatments. The origin of this specialization was not infectivity, which readily evolved and traded off weakly between the host species for both parasites. Instead, the overall specialization was caused by spore production, which did not evolve in any treatment. This suggests the existence of a strong trade-off between spore production in *A. franciscana* and spore production in *A. parthenogenetica*, making this trait a barrier to the evolution of generalism in this system. This study highlights that the shape of between-host trade-offs can be very heterogeneous across parasite traits, so that only some traits are pivotal to specialization.

## Introduction

Most parasites manifest a degree of specialization in nature, with niches that do not cover the entire community of potential hosts. This occurs even in communities of ecologically and physiologically similar host species (e.g. Antonovics et al. 2002, Hall et al. 2009, Streicker et al. 2013, Lievens et al. 2019), begging the question of why parasites do not evolve to extend their niche. Answering this question is particularly relevant when trying to predict the future evolution of a parasite, for example with regards to the emergence of new diseases (Cleaveland et al. 2001) or the impact of invasive hosts (Prenter et al. 2004, Kelly et al. 2009).

The evolution of host specialization – i.e. the evolution of parasitic niches – is generally studied through the lens of ecological specialization theory. A cornerstone of specialization theory is the assumption that adaptation to one environment trades off with adaptation to another (reviewed in e.g. Futuyma and Moreno 1988, Kassen 2002, Ravigné et al. 2009). The strength of the fitness trade-offs determines, to a large degree, whether specialist or generalist strategies evolve: strong trade-offs favor the evolution of specialists, while weak trade-offs favor the evolution of generalists. At first glance, therefore, we might view specialism in a parasite population as an indicator that there are strong fitness trade-offs between the hosts. This would imply that the parasite could never evolve to become a generalist. However, specialization is also governed by the availability and demography of the different environments (e.g. Bell and Reboud 1997, Ronce and Kirkpatrick 2001, Ravigné et al. 2009). Low encounter rates with the alternative host can maintain specialism in spite of weak trade-offs (Benmayor et al. 2009), especially if that host is an ecological sink (Holt and Gaines 1992, Holt and Hochberg 2002, Lenormand 2002, Ching et al. 2013). In this case, a demographic change in the host community could indeed prompt a shift towards generalism in the parasite. This scenario would seem especially likely among similar host species, where we might expect trade-offs to be weaker (cf. Hereford 2009). To disentangle the consequences of trade-offs from those of host availability, experimental evolution studies are necessary (Kassen 2002, Fry 2003).

Many experimental evolution studies have been done on host specialization, yielding a variety of outcomes: evidence for fitness trade-offs (e.g. Turner and Elena 2000, Yourth and Schmid-Hempel 2006, Agudelo-Romero et al. 2008, Legros and Koella 2010), mixed support for trade-offs (e.g. Agrawal 2000, Nidelet and Kaltz 2007, Magalhães et al. 2009, Bedhomme et al. 2012, Messina and Durham 2015, Meaden and Koskella 2017), selection for generalism when host availabilities fluctuate (e.g. Poullain et al. 2008, Legros and Koella 2010, Bedhomme et al. 2012, Magalhães et al. 2014), and complex effects of migration and host availability (e.g. Benmayor et al. 2009, Ching et al. 2013, Bono et al. 2015). However, very few experimental evolution studies take the natural context of parasite populations into account. Doing so can help disentangle the consequences of genetic constraints from the effects of host availability on the response to selection (see Jaenike and Dombeck 1998, Fellous et al. 2014). In addition, few studies look for the traits underlying fitness trade-offs. Parasite fitness is a composite of successful infection, host exploitation, and transmission. Some host specialization studies have shown that these traits can respond differently to selection on novel hosts (Magalhães et al. 2009, Bedhomme et al. 2012, Messina and Durham 2015), but their causal effects on parasite evolution have been largely unexplored (Hall et al. 2017). For example, low fitness in a new host may result from strong trade-offs in one or a few key traits, or from the accumulation of weak trade-offs in most traits. Only the former can prevent the evolution of a generalist parasite on the long term, so identifying whether such key traits occur is crucial to understanding host specialization.

In this study, we investigated whether trade-offs in host use limit the evolution of generalism in a natural host-parasite community, and if so, upon which traits these trade-offs act. We used the microsporidians *Anostracospora rigaudi* and *Enterocytospora artemiae* and their sympatric hosts, the brine shrimp *Artemia parthenogenetica* and *Artemia franciscana*. The two parasites are ecologically similar, can complete their life cycles on both hosts, and commonly infect both hosts in the field (Rode et al. 2013b). Nonetheless, they each show a degree of specialization in the lab: *A. rigaudi* has much higher fitness in *A. parthenogenetica*, while *E. artemiae*’s fitness is much higher in *A. franciscana* (Lievens et al. 2018). Furthermore, we have shown that *A. franciscana* is a sink host for *A. rigaudi* in the field, even though prevalences in this host can reach 100%, and that the same may be true for *A. parthenogenetica* and *E. artemiae* (Lievens et al. 2019). This combination of partial specialization and source-sink demography prompted us to disentangle host availability from trade-offs for these parasites. We manipulated their host environment by serially passaging them on one, the other, or an alternation of the two hosts. We then assayed the infectivity, virulence, and spore production rate of the evolved lines, and asked: [1] did manipulating the host environment affect the degree of specialization of the parasites?; [2] what was the role of the underlying traits?; and [3] were these results consistent with trade-offs in host use?

## Methods

### Hosts and parasites

#### Natural system

*Artemia* is a genus of small crustaceans occurring in hypersaline environments. Our study system, the saltern of Aigues-Mortes on the Mediterranean coast of France, contains two sympatric *Artemia* species. The first, *A. parthenogenetica*, is an asexual clade native to the area; the second, *A. franciscana*, is a sexual species that was introduced from North America in 1970 and has since become highly prevalent (Amat et al. 2005, Rode et al. 2013b). *A. parthenogenetica* is present from late spring to fall, while *A. franciscana* occurs year-round. When both species are present, they usually share the same microhabitats (Lievens et al. 2019).

The microsporidians *A. rigaudi* and *E. artemiae* are two of the most prominent parasites infecting *Artemia* in Aigues-Mortes. *A. rigaudi* is native to France; *E. artemiae* may be native or co-invasive with *A. franciscana* (Rode et al. 2013b). Both species are horizontally transmitted parasites of the gut epithelium. Infections continuously produce spores, which are released in the infected host’s faeces and ingested by new hosts while filter feeding (Rode et al. 2013a). This causes both survival and reproductive virulence (Rode et al. 2013b, Lievens et al. 2018), and there is little evidence for recovery (Lievens et al. unpublished data). In the field, prevalences of over 80% have been recorded for both microsporidians in both hosts, and coinfections are common (Lievens et al. 2019). Very little is known about coinfections, though there is circumstantial evidence that established infections can exclude new arrivals (Lievens et al. 2019).

Both *A. rigaudi* and *E. artemiae* are partially specialized: they can complete their life cycles on either host, but perform much better on one of the two (Lievens et al. 2018). *A. rigaudi*’s fitness is considerably higher in *A. parthenogenetica*, while *E. artemiae*’s is higher in *A. franciscana*. This is mainly caused by differences in spore production, although *E. artemiae* is also a poor infector of *A. parthenogenetica* and *A. rigaudi* exhibits higher-than-optimal virulence in *A. franciscana* (Lievens et al. 2018). The partial specialization is reflected in their field patterns: *A. rigaudi* cannot persist in nature if *A. parthenogenetica* is absent (Lievens et al. 2019). *A. rigaudi* epidemics therefore end in late fall and must be re-started every spring (possibly by cold-resistant spores, Lievens et al. unpublished data). It is unclear whether *E. artemiae* is equally dependent on *A. franciscana* in the field, but we suspect that it is (Lievens et al. 2019). In the lab, parasites can be maintained on either host.

#### Origin of experimental parasites

We obtained our experimental parasites from the same laboratory stocks of *A. rigaudi* and *E. artemiae* that were used by Lievens et al. (2019) to estimate infectivity, virulence, and spore production. The microsporidians in these stocks were collected in Aigues-Mortes and maintained on a mix of both hosts. Before starting the serial passages, we made sure that the microsporidian stocks were uncontaminated by using them to infect lab-bred hosts, testing those hosts for the presence of both microsporidians, and re-starting the stocks from singly infected hosts only (see Supplementary Material for more details). Note that although we tried to maximize the genetic diversity of our microsporidian stocks by using spores produced by both host species, originating in several sites and at different times, we do not know if the resulting populations were genetically diverse or not.

### Experimental evolution

We serially passaged the microsporidians *A. rigaudi* and *E. artemiae* on the host species *A. franciscana*, *A. parthenogenetica*, or an alternation of the two. After 10 passages, we assayed the infectivity, virulence, and spore production of each line, and compared these among treatments.

#### Experimental conditions

See Supplementary Methods.

#### Serial passages

We subjected *A. rigaudi* and *E. artemiae* to serial passaging under three evolutionary treatments: ‘*A. f*. host’, ‘*A. p.* host’, and ‘Alternating hosts’. In the first two regimes, the parasites encountered *only A. franciscana* or only *A. parthenogenetica*; in the third regime, the parasites encountered alternating passages of *A. franciscana* and *A. parthenogenetica*. Each microsporidian × treatment combination was replicated four times, producing a total of 24 parasite lines. Parasites underwent ten serial passages, each lasting three weeks. The protocol is depicted in Fig. 1; details can be found in the Supplementary Methods.

**Figure 1.**
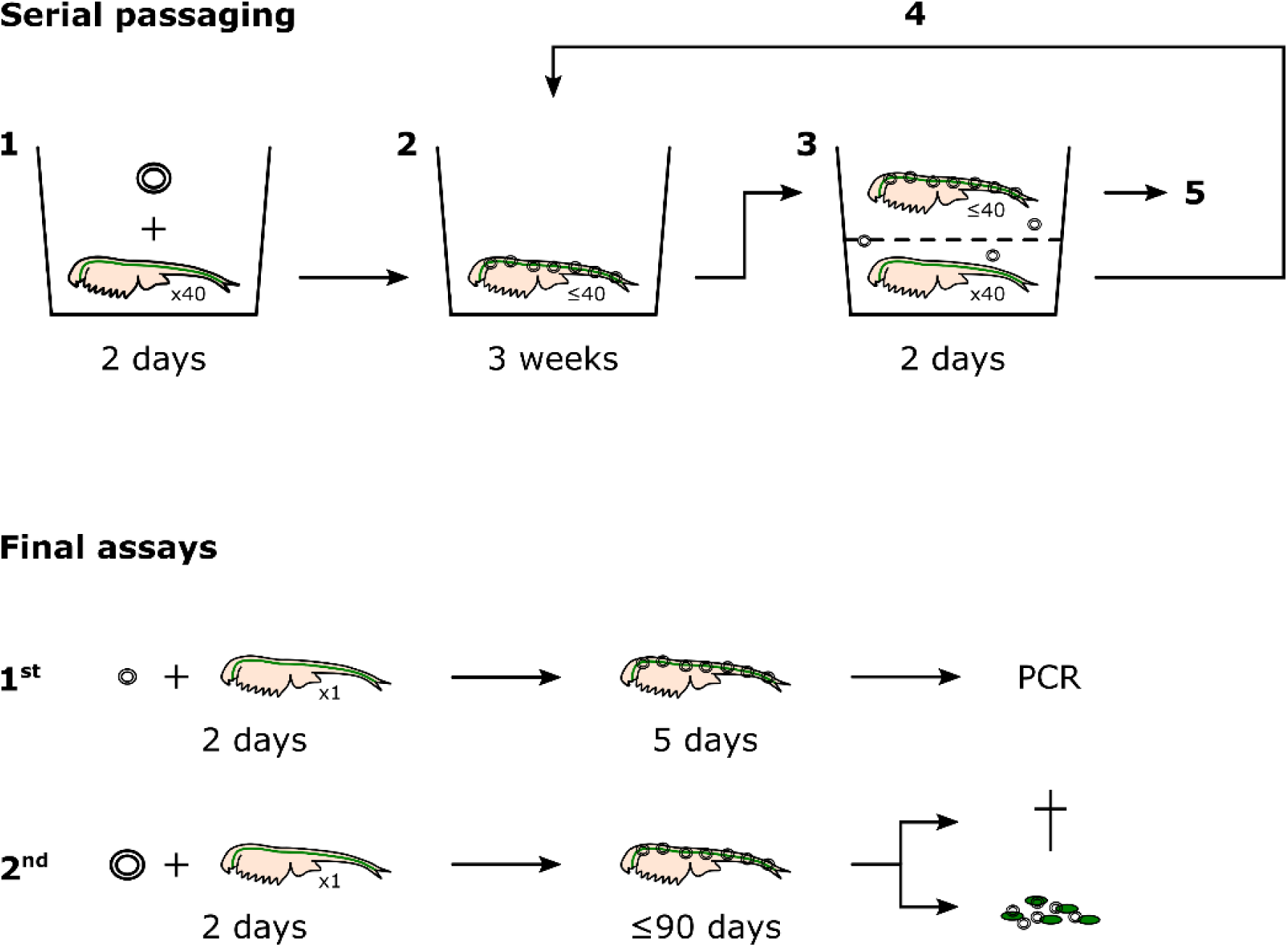
Experimental evolution protocol. **Serial passages:** [1] Passage 1 (P1): a group of 40 uninfected hosts was exposed to a saturating dose of stock *A. rigaudi* or *E. artemiae* spores, and [2] the infections were allowed to incubate. [3] Passaging (P1 → P2): the infection was transmitted naturally, by placing the surviving P1 hosts in a strainer above a new group of uninfected hosts. [4] The incubation and passaging steps were repeated for P2-P10. [5] After passaging, the surviving old hosts were counted (P1-P10) and used to estimate the population size of the parasite (P1, P4, P7), produce backup spore samples (P6), or produce the spores for the final assays (P10). **Final assays:** [1^st^] An uninfected host was exposed to a low dose of evolved *A. rigaudi* or *E. artemiae* spores, and PCR-tested for the presence of the microsporidian after a short incubation period. [2^nd^] An uninfected host was exposed to a saturating dose of evolved *A. rigaudi* or *E. artemiae* spores, and mortality and spore production were tracked for 90 days.

Two aspects of our passaging protocol should be pointed out: first, the time between passages (three weeks) is enough to allow infections to be transmitted within passaged groups (Rode et al. 2013a). Thus, low infection rates at the start of a passage could be compensated by high within-passage transmission. Second, we did not control the number of spores that were transmitted from one group of hosts to the next. In all passages after P1, the size of the inoculum depended on the parasite load of the old hosts. These two aspects meant that the lines were allowed to develop their own infection dynamics, just as they would in the field. Any stochasticity caused by variation in these demographic processes is explicitly included in the experiment.

#### Final assays

At the end of the serial passage experiment, we tested the infectivity, virulence, and spore production of each evolved line in both *A. franciscana* and *A. parthenogenetica*. We tested all surviving parasite lines based on the spores they produced at the end of P10. We also tested a subset of the parasite lines based on backup spores collected after P6, which we call the ‘revived’ lines (see Results and Table 1). Details on spore collection after P6 and P10 can be found in the Supplementary Methods.

**Table 1.**
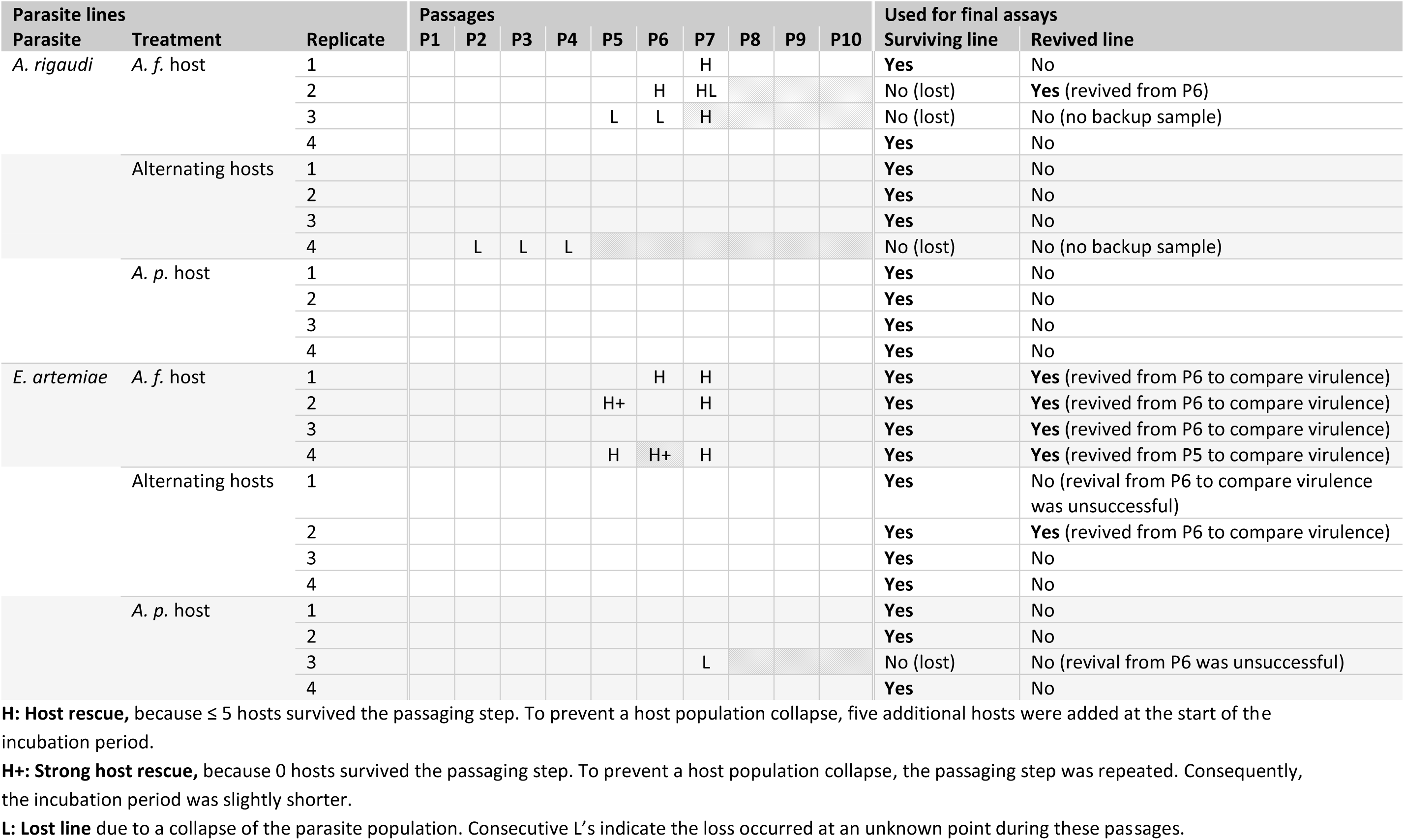
History of the parasite lines. Events that occurred during the serial passaging (abbreviated **H**, **H+**, and **L**) are noted at the relevant passaging step. For the final assays, we used all the surviving parasite lines, plus a set of lines that were revived from backup spore samples (see Results and Fig. 1).

We performed two final assays (described in Fig. 1, details in the Supplementary Methods). The first assay tested the infectivity of the evolved lines; it was replicated on 40 *A. parthenogenetica* and 40 *A. franciscana* individuals for each line. In parallel with the first, a second assay tested the virulence and spore production of each evolved line. This assay was also replicated on 40 *A. parthenogenetica* and 40 *A. franciscana* per evolved line. Because we suspected that there would be fewer infections when *A. parthenogenetica* was exposed to *E. artemiae*, we increased the level of replication for these combinations, adding an extra 20 *A. parthenogenetica*. We also included 80 control *A. parthenogenetica* and 80 control *A. franciscana*, which were not exposed to spores but otherwise treated identically. Spore production was measured by quantifying the number of spores produced by infected individuals over a two-day period after three weeks of infection; this corresponds to the window for transmission during the serial passage experiment.

### Statistical analyses

All analyses were carried out in R version 3.5.1 (R Core Team 2014), using the packages lme4 (linear mixed modeling, Bates et al. 2015) and survival (survival analyses, Therneau 2014). Unless stated otherwise, we built full models with the relevant experimental factors, and tested for the significance of effects using the likelihood ratio test. The specific models for each analysis are described below. If post-hoc testing was necessary, we used Tukey HSD tests from the packages multcomp (Hothorn et al. 2008) and lsmeans (Lenth 2016).

#### Serial passages

During the serial passage experiment, we collected data on host survival and parasite population size. Here, we tested whether these variables changed over the course of the experiment.

Host survival was quantified as the proportion of surviving hosts in each line at the end of each passage. Because we did not maintain “control” host populations during the serial passage experiment, host survival is relative (e.g. survival in ‘Alternating hosts’ vs. ‘*A. f.* host’ treatments), and can only be compared within host species; we therefore analyzed it separately for *A. franciscana* and *A. parthenogenetica*. Linear mixed models included survival as a binomial response variable, *Treatment*, *Passage number* (as a continuous variable measuring time), *Parasite species* and all interactions as fixed effects, and *Line* as a random effect. In addition, we included *Passage* as a random factor, to control for background variation in the quality of the hosts. Lines where parasites were lost (see Results and Table 1) were excluded.

To test whether the parasite population size changed, we built linear mixed models including *Treatment*, *Passage number* (as a continuous variable measuring time), and their interaction as fixed effects, and *Line* as a random variable. *A. rigaudi* and *E. artemiae* lines were analyzed separately. The population size was *ln*-transformed, and zero counts (lost lines) were excluded.

#### Final assays

In the final assays, we tested the effects of the passaging treatment on the infectivity, virulence, and spore production of the two parasites; we then compared a composite measure of parasite fitness. For each variable described below, analyses proceeded as follows. *A. rigaudi* and *E. artemiae* lines were analyzed separately. We began by testing whether surviving and revived lines were different, looking only at those treatments that included revived lines (models with fixed effects *Revival*, *Treatment, Assay host*, and their interactions). If they were not different, the revived lines were included in the subsequent analyses (models with fixed effects *Treatment*, *Assay host*, and their interaction). *Line* was always included as a random variable, or as a frailty variable for survival analyses.

Infectivity was analyzed as the proportion of infected individuals at the end of the first assay (a binomial response) in a generalized linear mixed model. For virulence, the effects of *Treatment* were tested using log-logistic survival models, stratified over *Assay host* (this allowed the host species to have a different baseline survival shape). So that the results could be interpreted in terms of survival relative to uninfected hosts, we included the survival data of the control hosts as an additional *Treatment* category. However, we excluded any hosts that had been exposed to a parasite but not infected (cf. Lievens et al. 2018)(see Results, Table 2 for the proportion of infected hosts). We also excluded any hosts that died before day 11 of the assay, because infection could not be reliably detected before this day (see Supplementary Methods). To analyze the effects on spore production, we used the spore count in the fecal sample as a negative binomial response variable in a generalized linear mixed model. Fecal samples were only pooled for infected individuals; uninfected hosts were therefore implicitly excluded from the model.

**Table 2.** Detection of infection in the second assay. Only hosts that died after day 10 are included here, to allow for the delay in detection time (see Supplementary Methods).

| Parasite species | Infected hosts |
| --- | --- |
| ***A. rigaudi*** |  |
| *A. franciscana* | 66 % |
| *A. parthenogenetica* | 70 % |
| ***E. artemiae*** |  |
| *A. franciscana* | 68 % |
| *A. parthenogenetica* | 34 % |

Finally, we used spore production and infectivity to produce a composite fitness measure for each line. We used a measure of fitness that was representative for the context of the experiment, being the projected number of infections occurring if the line were passaged onto a new set of susceptible hosts. We calculated this as the total number of spores produced by the surviving individuals over a two-day period after three weeks of incubation (thus virulence is implicit), multiplied by the infectiousness of a single spore. Infectiousness, the probability of a single spore to start an infection, was calculated based on the results of the first assay. Following an independent action model with birth-death processes, the infectiousness of one spore is − 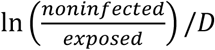, where *D* is the spore dose, in our case 750 spores (Schmid-Hempel 2011, pg. 225-6). We analyzed fitness using a linear mixed model, after *ln*+1 transformation.

## Results

### Serial passages

Of the twenty four parasite lines, four were lost during passaging due to a collapse of the parasite population (Table 1).

At the beginning of passages P5, P6, and P7, exceptionally high mortality occurred in several groups of new hosts as they were being exposed to the parasites produced by the old hosts (Table 1). These episodes were concentrated in the treatments ‘*A. f.* host’. To prevent the loss of these lines due to host population collapse, we added new hosts; if necessary, we repeated the passaging step (see Table 1). Notably, the passaging step from P5 to P6 was repeated three times without success for the line *E. artemiae* × ‘*A. f.* host’ – Replicate 4. The transfer was eventually achieved after 6 weeks of incubation in the P5 hosts, as the other lines were being passaged from P6 to P7. We denote this transfer as ‘P5 → P7’ for consistency, but P7 is only the 6^th^ passage for this particular line. To investigate whether these effects were due to increased virulence or demographic effects (increased parasite load), we included backup spores produced by these lines in the final assays (see below).

Host survival was not constant throughout the serial passage experiment, even when the background variation in host quality was taken into account (Fig. 2). As the passages progressed, the survival of *A. franciscana* in *A. rigaudi* × ‘*A. f.* host’ lines decreased as compared to the others (significant triple interaction, χ^2^(2) = 5.5, *p* = 0.02, Supp. Table 1; post-hoc −4.1 < *z* < −2.5, 0.0001 < *p* < 0.06). Because we could not separate the background host mortality from parasite-induced effects, we cannot say whether this change was due to increasing parasite-induced mortality in the *A. rigaudi* × ‘*A. f.* host’ lines, or to decreasing parasite-induced mortality in the other lines. For *A. parthenogenetica*, survival rates became progressively higher in *A. rigaudi* relative to *E. artemiae* lines, as well as in ‘Alternating hosts’ relative to ‘*A. p.* host’ lines (significant effects of *Parasite species* and *Treatment* in interaction with *Passage number*, χ^2^(1) = 11.4 and 8.6, *p* < 0.001 and *p* < 0.01, respectively, Supp. Table 1). Again, we could not distinguish between positive changes in *A. rigaudi* and ‘Alternating hosts’ lines or negative changes in *E. artemiae* and ‘*A. p.* host’ lines.

**Figure 2.**
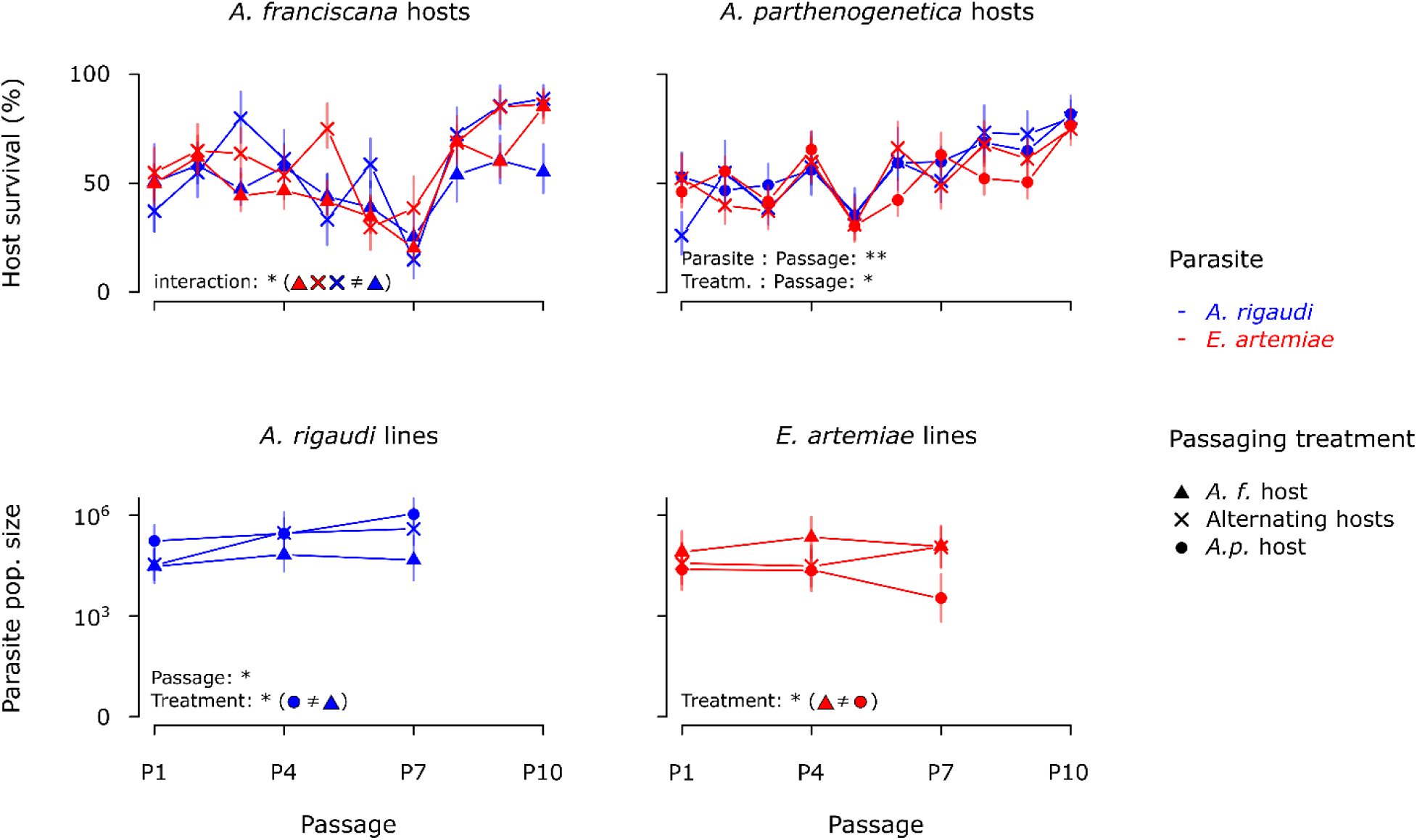
Host and parasite populations during the serial passaging. Host survival (top row) is a compound of background host mortality and parasite-induced mortality; parasite population size (bottom row) is the spore load in the surviving old hosts after passaging (*ln* scale). Lost lines (lines with a parasite population size of 0) are not included in the figure. The highest-order significant experimental variables are provided for each analysis (cf. Supp. Table 1); where relevant, the post-hoc results are shown in parentheses (see Results). Vertical bars represent the 95% CIs.

The estimated population size of the parasites also varied through time (Fig. 2). For *A. rigaudi*, the population grew over the course of the experiment (χ^2^(1) = 10.0, *p* < 0.01 for *Passage number*, Supp. Table 1), and was significantly larger for lines evolving on *A. parthenogenetica* than for lines evolving on *A. franciscana* (χ^2^(2) = 7.2, *p* = 0.03 for *Treatment*, Supp. Table 1; post-hoc *z* = 2.7, *p* = 0.02). For *E. artemiae*, only the passaging regime impacted the population size, which was significantly higher in lines evolving on *A. franciscana* than in those evolving on *A. parthenogenetica* (χ^2^(2) = 10.4, *p* < 0.01 for *Treatment*, Supp. Table 1; post-hoc *z* = 3.3, *p* < 0.01).

### Final assays

During the final assays, we tested all surviving evolved lines, as well as a set of lines revived from the backup P6 spore samples (Table 1). These included all the lines in the combination *E. artemiae* × ‘*A. f.* host’, most of which experienced a period of exceptional mortality during the transmission events before the end of P6 (Table 1). The two *E. artemiae* × ‘Alternating hosts’ lines whose P6 hosts were *A. franciscana* (Replicates 1 & 2) were also revived to act as controls for the effect of storage, but revival was only successful for Replicate 2. Finally, we succeeded in reviving the spores of the lost line *A. rigaudi* × ‘A. f. host’ – Replicate 2.

In the first assay, we tested for effects of passaging treatment on infectivity (Fig. 3, replicates shown in Supp. Fig. 1). The infectivity of *A. rigaudi* was unaffected by storage effects (χ^2^(1) = 1.6, *p* = 0.21), and did not change in response to passaging treatment (χ^2^(2) = 2.8, *p* = 0.25, Supp. Table 2); it tended to be higher in *A. parthenogenetica* (χ^2^(1) = 3.7, *p* = 0.054, Supp. Table 2). In contrast, the infectivity of *E. artemiae* was reduced by storage at 4°C (χ^2^(1) = 4.6, *p* = 0.03. dashed lines in Supp. Fig. 1), so the revived lines were excluded from further analysis. The infectivity of surviving *E. artemiae* lines was generally higher in *A. franciscana* than in *A. parthenogenetica*, but the difference was less strong after passaging on ‘Alternating hosts’ and ‘*A. p.* host’ (χ^2^(2) = 8.1, *p* = 0.02 for interaction effect, Supp. Table 2).

**Figure 3.**
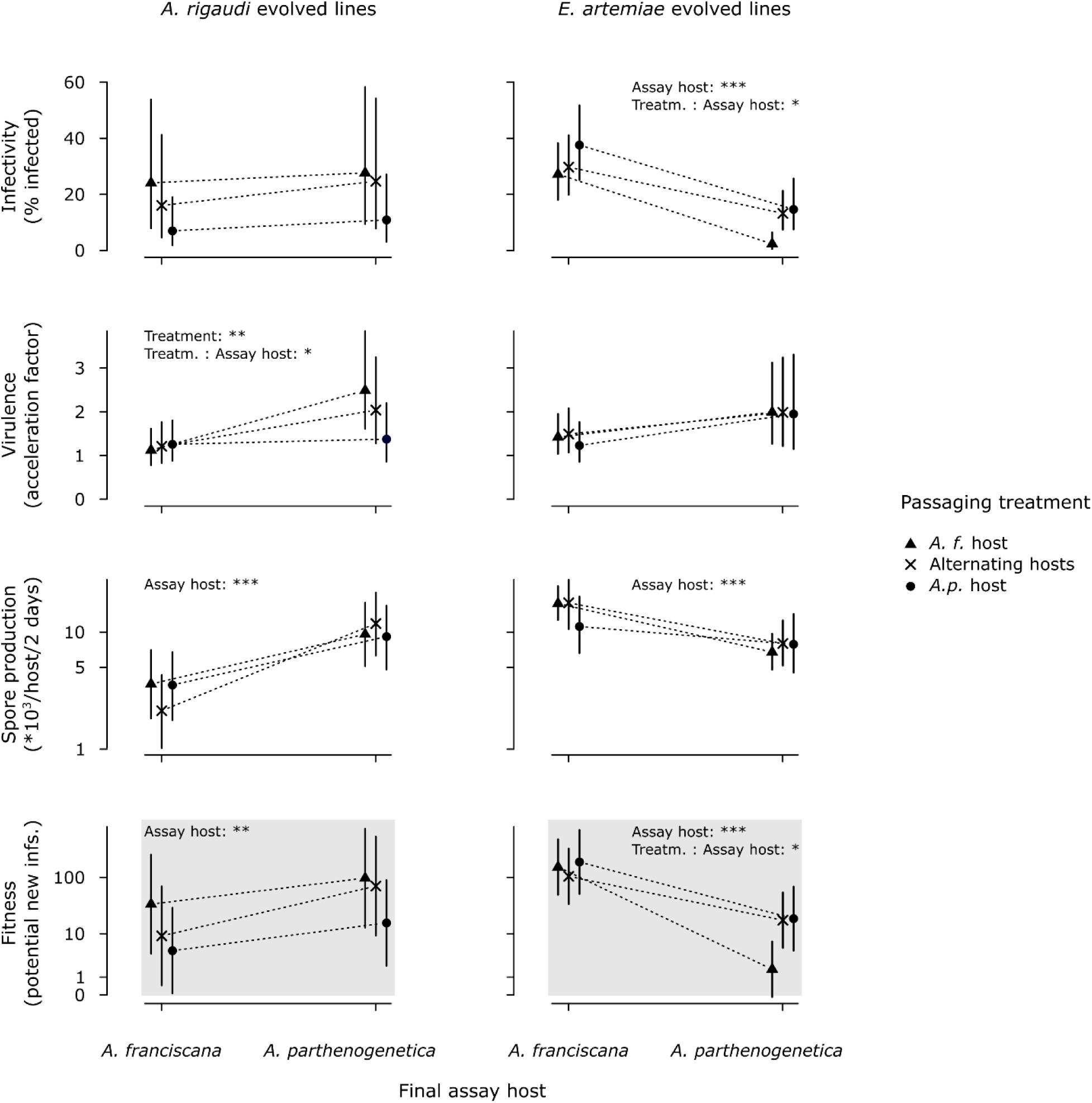
Results of the final assays. Infectivity, virulence, and spore production were measured directly; fitness was calculated based on these (differentiated by the gray background). Infectivity is the percentage of hosts infected during the first assay. Virulence and spore production were measured in the second assay: virulence is the acceleration factor (the ratio of time-until-death) compared to the unexposed controls of the same species, spore production is the number of spores produced per (surviving) infected host at the time of passaging (*ln* scale). Fitness is the projected number of infections at passaging (*ln* + 1 scale). The significant experimental variables are noted for each microsporidian × treatment combination (except that of *Assay host* for virulence, see Supp. Table 2). Vertical bars represent the 95% CIs.

In the second assay, we tested for effects of passaging treatment on virulence and spore production (Fig. 3, replicates shown in Supp. Fig. 2 and 3). As expected, we detected infection in the majority of the exposed hosts in all host-parasite combinations except A. parthenogenetica-E. artemiae (Table 4). No infection was detected for the line *A. rigaudi* × ‘*A. p.* host’ – Replicate 1, so it was excluded from further analyses.

For both *A. rigaudi* and *E. artemiae*, parasite-induced mortality was unaffected by storage at 4°C (respectively χ^2^(1.0) = 0.2, *p* = 0.64 and χ^2^(1.2) = 0.9, *p* = 0.39). The revived lines were therefore included in the analyses. Overall, mortality was higher for *A. parthenogenetica*. For *A. rigaudi*, there was an additional effect of passaging treatment: when assayed on *A. parthenogenetica*, virulence was highest for lines passaged on *A. franciscana*, intermediate for lines passaged on alternating hosts, and lowest for lines passaged on *A. parthenogenetica* itself (χ^2^(3.0) = 10.4, *p* = 0.02 for interaction effect, Supp. Table 2; Fig. 3, replicates shown in Supp. Fig. 2). For *E. artemiae* lines, background mortality was also higher for *A. parthenogenetica* (χ^2^(4) = 58.1, *p* < 0.0001, Supp. Table 2), but infected hosts did not die faster than unexposed hosts: virulence was not affected by passaging treatment, nor by the interaction between treatment and assay host (χ^2^(3.6) = 3.7, *p* = 0.38 and χ^2^(4.5) = 5.8, *p* = 0.27, respectively, Supp. Table 2; Fig. 3, replicates shown in Supp. Fig. 2).

Similarly, spore production at passaging was unaffected by storage at 4°C (χ^2^(1) = 0.2, *p* = 0.69 for *A. rigaudi*; χ^2^(1) = 0.8, *p* = 0.38 for *E. artemiae*), so all lines were included in the further analyses. Spore production was higher in *A. parthenogenetica* for *A. rigaudi* and in *A. franciscana* for *E. artemiae* (χ^2^(1) = 14.8 and = 16.5, *p* = 0.0001 and < 0.0001, respectively, Supp. Table 2; Fig. 3, replicates shown in Supp. Fig. 3). However, there were no effects of treatment, nor of the interaction between treatment and assay host (χ^2^(2) ≤ 0.7 and ≤ 1.7, *p* ≥ 0.71 and ≥ 0.43, respectively, Supp. Table 2; Fig. 3, replicates shown in Supp. Fig. 3).

Finally, we analyzed an overall fitness measure for each line: the projected number of hosts that would be infected at passaging (gray panels in Fig. 3). For *A. rigaudi*, storage at 4°C had no effect on the composite traits of fitness (see above), so the single revived line was included in the analysis. We also included the line *A. rigaudi* × ‘*A. p.* host’ – Replicate 1, which failed to infect hosts in the second assay, with fitness set to 0 (excluding the line did not change the results). *A. rigaudi* fitness was always higher when tested on *A. parthenogenetica*, with no effect of passaging treatment, or of the interaction between treatment and assay host (χ^2^(1) = 11.9, χ^2^(2) = 2.2 and 2.5, *p* < 0.001, = 0.34 and = 0.29, respectively, Supp. Table 2; gray panels in Fig. 3). For *E. artemiae*, in contrast, the patterns of fitness mirrored those of infectivity. As storage at 4°C affected infectivity (see above), the revived lines were excluded. *E. artemiae* fitness was always lower in *A. parthenogenetica*, but less so after passaging on ‘Alternating hosts’ and ‘*A. p.* host’ (χ^2^(2) = 6.4, *p* = 0.04 for interaction effect, Supp. Table 2; gray panels in Fig. 3).

## Discussion

We investigated the evolution of host specialization, and its underlying traits, in the microsporidian parasites *A. rigaudi* and *E. artemiae*. In the field, these parasites infect two sympatric species of *Artemia*, each with a degree of host specialization: *A. rigaudi* is preferentially adapted to *A. parthenogenetica*, and *E. artemiae* to *A. franciscana* (the “matched” hosts, Lievens et al. 2018). To test whether this pattern is shaped by host availability or by fitness trade-offs, we experimentally evolved the parasites on one or both of their natural hosts. We found that the parasites remained partially specialized in all passaging conditions. The different parasite traits did not play an equal role in this outcome: spore production remained specialized in both parasites, infectivity showed a generalist pattern in *A. rigaudi* and readily evolved towards generalism in *E. artemiae*, and virulence played a minor role. Our results are consistent with a strong trade-off acting on spore production and a weak trade-off on infectivity, and suggest that spore production is the key trait preventing the evolution of generalism in this system.

### The evolution of specialization and its underlying traits

Our first conclusion is that both *A. rigaudi* and *E. artemiae* display a robust pattern of specialization: the fitness of both microsporidians was higher in the matched hosts than in the mismatched hosts, even after extended passaging on the latter (Fig. 3, gray panels). This result is consistent with our previous ecology- and life history-based findings (Lievens et al. 2018, 2019).

*A. rigaudi*’s specialization for *A. parthenogenetica* was caused by a disparity in spore production. This parasite produced many more spores in *A. parthenogenetica*. Neither infectivity nor spore production changed detectably during the serial passages, but the passaging treatment did affect virulence (Fig. 3). When tested in *A. parthenogenetica*, *A. rigaudi* lines that had evolved on that host were less virulent than lines that had evolved on *A. franciscana*. Whether this was due to an incidentally high virulence on a ‘novel’ host, or to an adaptive decrease in virulence on a ‘known’ host, is unknown, but both are plausible (Woolhouse et al. 2001, Alizon et al. 2009). The effect of virulence on overall fitness was minor, however, so the parasite stayed equally specialized for *A. parthenogenetica* in all treatments (Fig. 3, gray panels).

For *E. artemiae*, specialization was apparent for spore production and infectivity. *E. artemiae* spores had a higher chance of infecting *A. franciscana*, and *E. artemiae* infections also produced more spores in *A. franciscana* (Fig. 3). Compounded, these two traits produce a clear pattern of specialization (Fig. 3, gray panels). Unlike that of *A. rigaudi*, however, *E. artemiae*’s fitness did evolve in some treatments. *E. artemiae* lines whose passaging history included *A. parthenogenetica* had a higher fitness on this host, while their fitness in *A. franciscana* was not detectably changed (compare cross & circle to triangle in Fig. 3). *E. artemiae* can thus evolve a more generalist strategy without a detectable trade-off. This observation supports the mounting evidence that “costs” of adaptation to different environments may not always be present, as expected theoretically (Fry 1996, Lenormand et al. 2018) and observed empirically (Falconer 1990, Agrawal 2000, Kassen 2002, Nidelet and Kaltz 2007, Magalhães et al. 2009, Bedhomme et al. 2012, Remold 2012, Gallet et al. 2014, Messina and Durham 2015). *E. artemiae*’s fitness change was driven by a change in infectivity, while virulence and spore production were static. Interestingly, changes in infectivity have also been found to drive the evolution of specialists and generalists in the microsporidian *Brachiola algerae*, although in this case there was a correlated loss of infectivity in other hosts (Legros and Koella 2010).

The difference in infectivity among the evolved lines of *E. artemiae* can be interpreted in two ways: its infectivity in *A. parthenogenetica* either decreased when the parasite was no longer exposed to this host, or increased when the parasite was forced to persist in it. We consider the second to be more likely. Overall, evolution is likely to have occurred from standing genetic variation, because we tried to maximize the diversity of our microsporidian stocks and used large inoculum sizes. However, *de novo* mutations may also have arisen due to the large population sizes (Fig. 2) and appreciable duration of the experiment (> 40x the time necessary for a detectable intra-host population to accumulate, Rode et al. 2013a). A decrease of infectivity when *E. artemiae* was not exposed to *A. parthenogenetica* could have been achieved by an increase in frequency of conditionally deleterious genotypes - neutral in *A. franciscana* and deleterious in *A. parthenogenetica* (de novo, Kawecki 1994, or by drift, Yourth and Schmid-Hempel 2006). However, given the large population sizes during serial passaging (Fig. 2), we doubt that such processes occurred. It is more likely that passaging on *A. parthenogenetica* caused genotypes that were better suited to this species to accumulate. This hypothesis is also supported by previous experimental results, which describe the infectivity of the stock population of *E. artemiae* as resembling that of the ‘*A. f.* host’ evolved lines (Lievens et al. 2018). If so, adaptation likely occurred through an increase in frequency of genotypes that were beneficial in *A. parthenogenetica* and neutral in *A. franciscana*. Another possibility is that adaptation occurred in all passaging treatments, but that adaptation to *A. parthenogenetica* had an incidental positive effect in *A. franciscana* that matched the adaptation to the ‘*A. f.* host’ treatment.

*E. artemiae*’s virulence did not differ among treatments at the end of the serial passaging, and we found no evidence that it evolved over the course of the experiment. In particular, we found no evidence that the high death rates caused by *E. artemiae* in *A. franciscana* between P4 and P6 were caused by a higher virulence (Supp. Fig. 2). Instead, demographic effects were the likely culprit: *E. artemiae*’s spore production in *A. franciscana* does not plateau at a certain maximum (Lievens et al. unpublished data), so a higher infective dose in this combination might lead to a higher transmission rate, which would increase the infective dose and so on, until the number of invading spores was so high that recipient hosts were overwhelmed (e.g. Ebert et al. 2000).

### Trade-offs in infectivity and spore production

An important advantage of this study is that *A. rigaudi* and *E. artemiae* share the same context: the two parasites are ecologically similar, sympatric, and infect the same host species, so we can reasonably expect that they are subject to similar life history and environmental constraints. Below, we take advantage of this to compare the evolved changes in infectivity and spore production for the two microsporidians, arriving at the compelling conclusion that the strength of their life history trade-offs is trait-dependent, but that the traits respond similarly in the two species.

The observed changes in infectivity can be explained by the existence of a weak trade-off between infectivity in *A. franciscana* and *A. parthenogenetica*. Consider first *E. artemiae*, whose ability to infect *A. parthenogenetica* improved when passaged on that host, without attendant losses in *A. franciscana*. Such cost-free adaptation could arise if the ancestral ‘*A. franciscana*-adapted’ infectivity of *E. artemiae* was located slightly below the boundary of a weak trade-off curve (such as would be expected if the ancestral population was not perfectly adapted to the conditions of the experiment, Fry 2003). There would then be little improvement possible in *E. artemiae*’s fitness on *A. franciscana*, but a substantial improvement in *A. parthenogenetica* could easily be achieved (blue arrow in Fig. 4)(Martin and Lenormand 2015), as seen when *E. artemiae* was passaged on this host (Fig. 3). The infectivity of *A. rigaudi* can be interpreted in the same context. *A. rigaudi*’s ancestral infectivity is largely generalist (Lievens et al. 2018), so that the potential improvements in fitness would be small in either direction, thus producing the unchanged infectivity that we observed across treatments (Fig. 3).

**Figure 4.**
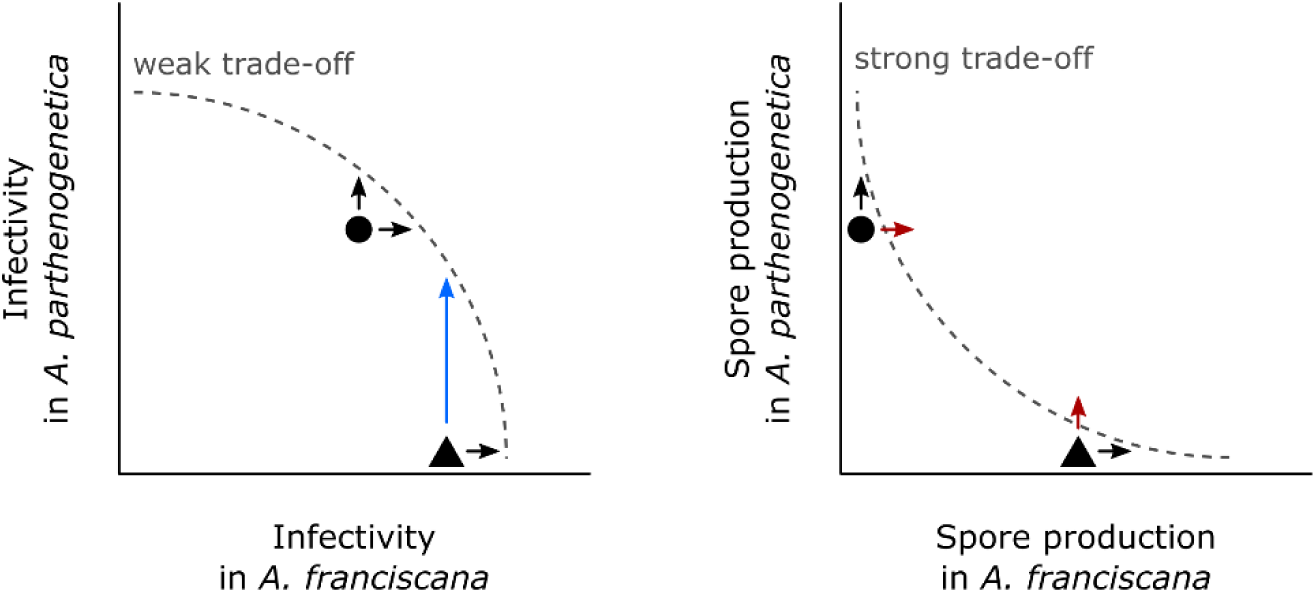
We speculate that the observed patterns of specialization in infectivity and spore production are determined by weak and strong trade-offs in performance between *A. franciscana* and *A. parthenogenetica* (see text for more information). Symbols: *A. rigaudi*, circle; *E. artemiae*; triangle. Black arrows, small changes are possible; blue arrow, a large change is possible; red arrows, changes in the direction of selection are not possible unless preceded by reverse specialization.

The weak trade-off model implies that the evolution of generalist infectivity should be straightforward, begging the question of why *E. artemiae*’s ancestral population remained specialized for this trait. We speculate that the specialization is maintained by source-sink dynamics in the natural host-parasite community. In the field, *E. artemiae* is present year-round. *A. parthenogenetica* hosts are only present from late spring to fall, so the parasite population predominantly infects, and evolves on, *A. franciscana* (Lievens et al. 2019). In this case, adaptations towards increased infectivity in the mismatched host may be continually eroded by selection in the matched host (Holt and Hochberg 2002, Lenormand 2002). By forcing *E. artemiae* to evolve on *A. parthenogenetica*, we blocked these source-sink dynamics, allowing generalist infectivity to evolve. In comparison, *A. rigaudi* almost exclusively occur in communities containing both host species (Lievens et al. 2019), potentially explaining why this microsporidian had already evolved generalist infectivity. Of course, other factors than demography could affect the evolution of infectivity in the field, including trade-offs with other traits (Alizon and Michalakis 2015) and competition between parasites (Mideo 2009).

The second important trait for *A. rigaudi* and *E. artemiae* was spore production, which remained strongly specialized in all treatments (Fig. 3). This unresponsiveness to the evolutionary treatment could have three explanations. First, high stochastic variation in our experimental design (drift and measurement error) could deprive us of the power to detect any adaptive change. The increased infectivity observed for *E. artemiae* makes this explanation unconvincing for any trait under similar levels of selection. As spore production is directly related to parasite fitness, there is no reason to expect weaker selection on this trait compared to infectivity. Second, there could be a complete lack of genetic diversity in this trait – either in the initial inocula or due to *de novo* mutations. This explanation is also unlikely given the way we assembled our initial inoculum (see above) and the observation of a genetic response in other traits (virulence in *A. rigaudi* and infectivity in *E. artemiae*). The populations we used were not generally devoid of genetic variation, and there is no reason to expect that mutation rates are inherently lower for spore production. We also observed phenotypic variation for spore production trait among lines (Supp. Fig. 3), which reinforces this point. The third explanation, which we decidedly favor, is that there is a strong trade-off between spore production in *A. franciscana* and spore production in *A. parthenogenetica*. Such a trade-off would allow small improvements in the direction of increased specialization (black arrows in Fig. 4), but make improvements on the novel host much more difficult to achieve (red arrows in Fig. 4), thereby preventing the emergence of more generalist phenotypes. Mechanistically, a strong trade-off could be related to the distinct strategies of host exploitation necessary to thrive in *A. franciscana* and *A. parthenogenetica*. The precise physiology of the host species is likely to be different (they have been diverging for an estimated 40 million years, Baxevanis et al. 2006), and indeed the mechanisms of virulence and within-host regulation employed by *A. rigaudi* and *E. artemiae* in their matched hosts differ (Lievens et al. 2018, Lievens et al. unpublished data), with *A. rigaudi* causing more survival virulence, and *E. artemiae* more reproductive virulence. Successful exploitation of *A. franciscana* and *A. parthenogenetica* could therefore require very different toolkits, preventing the evolution of generalism and reducing the likelihood of a host switch (cf. Gemmill et al. 2000).

Taken together, our results provide strong evidence that the microsporidians’ traits are constrained by different trade-off shapes. Intriguingly, while the trade-offs are trait-specific, they are not species-specific. It seems that while *A. franciscana* and *A. parthenogenetica* are not physiologically similar enough to allow the evolution of generalism, *A. rigaudi* and *E. artemiae* are ecologically similar enough to share the same constraints.

### Perspectives

Overall, we find that the natural specialization of *A. rigaudi* and *E. artemiae* is primarily shaped by a strong trade-off acting on spore production. Spore production is therefore a key trait blocking the evolution of generalism. However, host availability did affect the degree of specialization, by allowing the evolution of generalist infectivity. We therefore predict that the natural population of *E. artemiae* may eventually evolve to become less specific, as *A. rigaudi* is, but that neither parasite is likely to become a true generalist or to switch hosts.

It is worth noting that our conclusions would have been very different if we had not measured the parasites’ traits separately. Based on the overall fitness (gray panels in Fig. 3), we would have concluded that *A. rigaudi* was unable to adapt to its mismatched host, while *E. artemiae* was able to evolve towards generalism after exposure to *A. parthenogenetica*. This would have suggested that the two parasites had asymmetrical fitness trade-offs between hosts: a strong trade-off for *A. rigaudi*, and a weaker trade-off for *E. artemiae*. Ignoring the individual traits, therefore, can have important consequences for the interpretation of field patterns, and for the prediction of a parasite’s future evolution. Our results suggest that more theoretical studies of specialization should be set in a multi-trait context, with each trait able to exhibit weak or strong trade-offs and evolve accordingly. Such studies would be better equipped to describe the continuum between generalist and specialist strategies, and to single out the traits favoring their evolution.

## Acknowledgements

We thank C. Gilliot for her help establishing the stock populations, and C. Gilliot and T. Mathieu for their help designing the experimental equipment. We are also grateful to R. Zahab for her assistance with PCRs, and to R. Zahab, L. Olazcuaga, and J. Pantel for helping out in times of need. E. J. P. L. received financial support from a French Ministry of Research fellowship. TL and YM acknowledge support from CNRS and IRD.

Version 4 of this preprint has been peer-reviewed and recommended by Peer Community In Evolutionary Biology (https://doi.org/10.24072/pci.evolbiol.100090).

## Data accessibility

Data and analyses have been uploaded to Zenodo, doi 10.5281/zenodo.3476544.

