## Supplemental material for "Trait-specific trade-offs prevent niche expansion in two parasites"

#### SUPPLEMENTARY MATERIAL

**Supplementary Table 1.** Significance of the experimental variables for host mortality and parasite population size during the serial passages. *Treatment* refers to the passaging treatments '*A. f.* host', '*A. p.* host' and 'Alternating hosts'. All models included parasite line as a random effect; host mortality models also contained *Passage* as a random factor. Significance was tested using the likelihood ratio test; if there were significant interactions, the independent contributions of the effects were tested separately using Type III sum of squares (Wald $\chi^2$ test; in italics).

| Response variable | Fixed effect | df | $\chi^2$ | p |
|---|---|---|---|---|
| **Changes in host mortality** | | | | |
| *A. franciscana* | Treatment | *1* | *2.7* | *0.10* |
| | Parasite species | *1* | *1.8* | *0.18* |
| | Passage number | *1* | *0.0* | *0.99* |
| | Treatment : Parasite species | *1* | *3.1* | *0.08* |
| | Passage number : Treatment | *1* | *16.9* | *< 0.0001* |
| | Passage number : Parasite species | *1* | *8.4* | *< 0.01* |
| | Passage number : Treatment : Parasite species | 2 | 5.5 | 0.02 |
| *A. parthenogenetica* | Treatment | *1* | *6.6* | *0.01* |
| | Parasite species | *1* | *3.1* | *0.08* |
| | Passage number | *1* | *22.7* | *< 0.0001* |
| | Treatment : Parasite species | 1 | 1.4 | 0.23 |
| | Passage number : Treatment | 1 | 8.6 | < 0.01 |
| | Passage number : Parasite species | 1 | 11.4 | < 0.001 |
| | Passage number : Treatment : Parasite species | 2 | 0.4 | 0.54 |
| **Changes in parasite population size** | | | | |
| *A. rigaudi* | Treatment | 2 | 7.2 | 0.03 |
| | Passage number | 1 | 10.0 | < 0.01 |
| | Passage number : Treatment | 2 | 2.2 | 0.33 |
| *E. artemiae* | Treatment | 2 | 10.4 | < 0.01 |
| | Passage number | 1 | 0.0 | 0.95 |
| | Passage number : Treatment | 2 | 3.9 | 0.14 |

**Supplementary Table 2.** Significance of the experimental variables for fitness, infectivity, virulence, and spore production during the final assay. *Treatment* refers to the passaging treatments '*A. f.* host', '*A. p.* host' and 'Alternating hosts'. For virulence, *Treatment* also included a 'Control' category of unexposed hosts, so that in these analyses the effect of *Assay host* reflects the difference in background mortality (note that in Fig. 4, the effect of background mortality is removed by representing virulence as the survival of infected hosts compared to that of controls of the same *Assay host*). All models included evolved line as a random or frailty effect. Significance was tested using the likelihood ratio test; if there were significant interactions, the independent contributions of the effects were tested separately using Type III sum of squares (Wald $\chi^2$ test; in italics). Significant type III sum of squares results are not represented in Fig. 4.

| Response variable | Fixed effect | df | $\chi^2$ | p |
|---|---|---|---|---|
| **Infectivity** | | | | |
| *A. rigaudi* | Treatment | 2 | 2.8 | 0.25 |
| | Assay host | 1 | 3.7 | 0.05 |
| (revived line included) | Treatment : Assay host | 2 | 0.6 | 0.73 |
| *E. artemiae* | Treatment | *2* | *1.8* | *0.41* |
| | Assay host | *1* | *19.7* | *< 0.0001* |
| (revived lines excluded) | Treatment : Assay host | 2 | 8.1 | 0.02 |
| **Virulence** | | | | |
| *A. rigaudi* | Treatment | *3* | *15.8* | *0.001* |
| | Assay host | *1* | *4.9* | *0.03* |
| (revived line included) | Treatment : Assay host | 3.0 | 10.4 | 0.02 |
| *E. artemiae* | Treatment | 3.6 | 3.7 | 0.38 |
| | Assay host | 4.0 | 58.1 | < 0.0001 |
| (revived lines included) | Treatment : Assay host | 4.5 | 5.8 | 0.27 |
| **Spore production** | | | | |
| *A. rigaudi* | Treatment | 2 | 0.0 | 0.99 |
| | Assay host | 1 | 14.8 | 0.0001 |
| (revived line included) | Treatment : Assay host | 2 | 2.1 | 0.36 |
| *E. artemiae* | Treatment | 2 | 0.7 | 0.71 |
| | Assay host | 1 | 16.0 | < 0.0001 |
| (revived lines included) | Treatment : Assay host | 2 | 1.7 | 0.43 |
| **Fitness** | | | | |
| *A. rigaudi* | Treatment | 2 | 2.2 | 0.34 |
| | Assay host | 1 | 11.6 | < 0.001 |
| (revived line included) | Treatment : Assay host | 2 | 2.5 | 0.29 |
| *E. artemiae* | Treatment | *2* | *0.4* | *0.83* |
| | Assay host | *1* | *35.5* | *< 0.0001* |
| (revived lines excluded) | Treatment : Assay host | 2 | 6.4 | 0.04 |

**Supplementary Figure 1.** Infectivity of the evolved lines during the final assays. Top: *A. rigaudi* lines; bottom: *E. artemiae* lines. Solid points and lines represent surviving lines (at P10), while hollow points, '+' signs and dashed lines indicate revived lines (P6 backup spores used); points are connected per parasite line. The predicted infectivity is shown in black, with the vertical bar indicating the 95% CI (analyses were done separately for each parasite and did not include the revived *E. artemiae* lines).

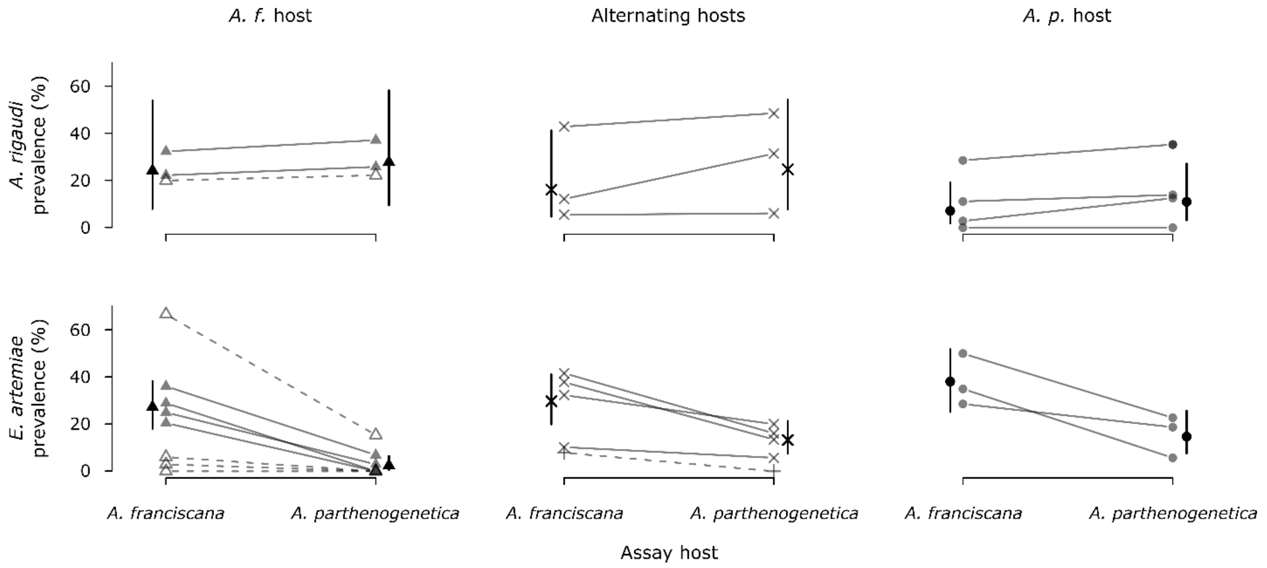

**Supplementary Figure 2.** Survival of infected hosts during the final assays (from day 11 onwards). Virulence is expressed by the acceleration factor (the ratio of time-until-death) of each line compared to the unexposed controls of the same species. Top: *A. rigaudi* lines; bottom: *E. artemiae* lines. Solid points and lines represent surviving lines (at P10), while hollow points, '+' signs and dashed lines indicate revived lines (P6 backup spores used); points are connected per parasite line. The predicted virulence is shown in black, with the vertical bar indicating the 95% CI (analyses were done separately for each parasite).

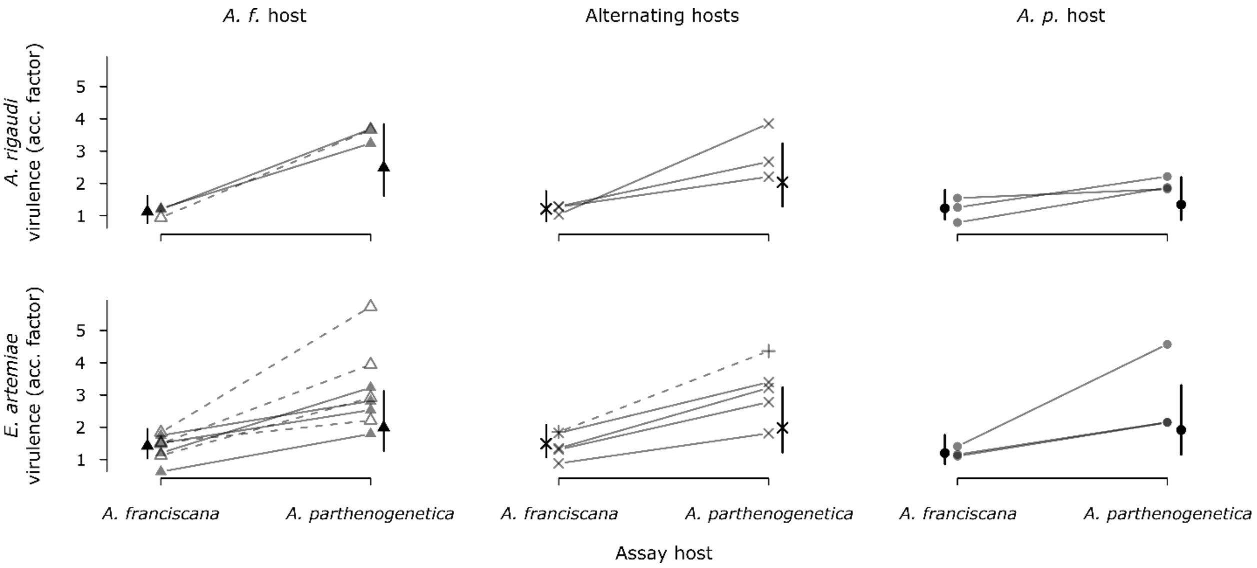

**Supplementary Figure 3.** Spore production during the final assays. Shown is the number of spores produced per (surviving) infected host after three weeks of incubation, over a two-day period. Top: *A. rigaudi* lines; bottom: *E. artemiae* lines. Solid points and lines represent surviving lines (at P10), while hollow points, '+' signs and dashed lines indicate revived lines (P6 backup spores used); points are connected per parasite line. The predicted spore production is shown in black, with the vertical bar indicating the 95% CI (analyses were done separately for each parasite).

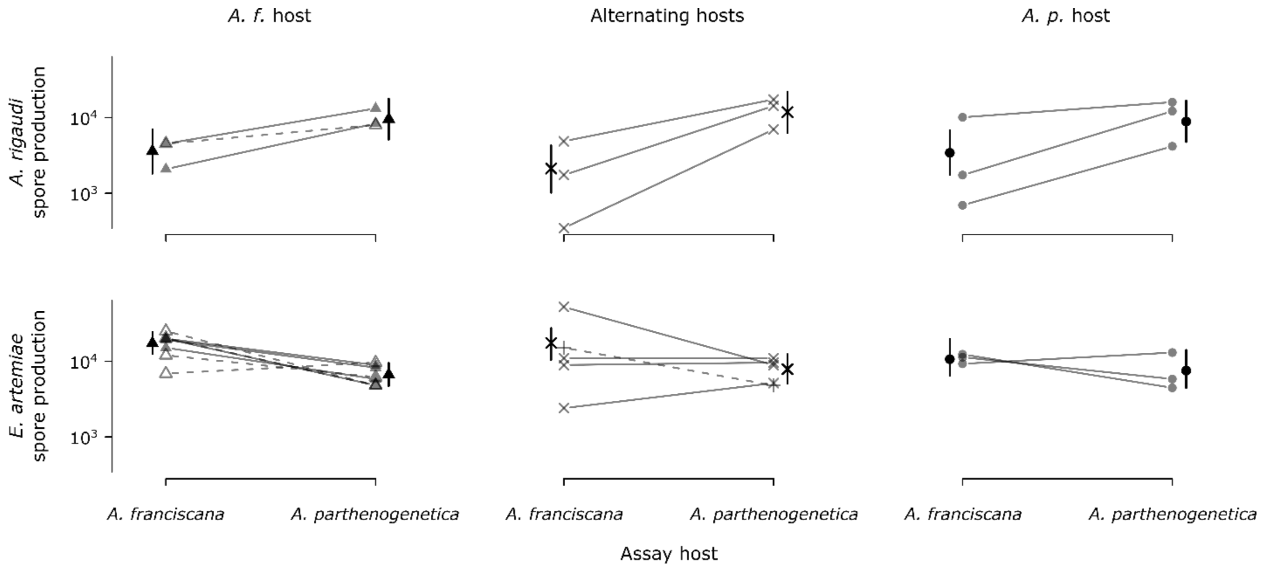

### SUPPLEMENTARY METHODS

#### Hosts and parasites

##### *Origin of experimental parasites*

We used the field-based laboratory stocks of *A. rigaudi* and *E. artemiae* used by Lievens et al. (2018), who describe them as follows:

> "We created stocks of *A. rigaudi* and *E. artemiae* for use in the experiment by combining infected *Artemia* from various sites in Aigues-Mortes between October 2014 and March 2015. We added new infected hosts to the stocks whenever we found field populations that were heavily infected with either *A. rigaudi* or *E. artemiae*. We also regularly added uninfected, lab-bred *Artemia* to help maintain the infection. We selected both infected *A. franciscana* and infected *A. parthenogenetica* from the field, and maintained each stock population on a mix of *A. franciscana* and *A. parthenogenetica* hosts ($n_{hosts}$ at any given time = ~20-~50 per microsporidian species). Thus, our stocks contained a mix of spores from different field sites and times, collected from and propagated on both host species."

Our field-based laboratory stocks of *A. rigaudi* and *E. artemiae* were susceptible to contamination with respectively *E. artemiae* and *A. rigaudi*, because we could not eliminate the possibility that a coinfected host had been added accidentally. To ensure we were using an inoculum containing only *A. rigaudi* or *E. artemiae*, we purified our stocks in three steps:

**1)** We infected lab-bred *Artemia* with spore samples taken from the field-based stock populations at different times. For *A. rigaudi*, we used spores sampled from the stock population on 28/11/2014, 5/12/2014 and 13/2/2015. For *E. artemiae*, we used spores sampled from the stock population on 13/2/2015 and 25/2/2015. Each of these samples was taken by collecting the feces produced by the stock hosts during one week, and had been stored at 4°C since. For each of the five spore solutions, we infected 16 large, adult *A. parthenogenetica* and 32 medium-sized, juvenile *A. franciscana*. *A. parthenogenetica* were isolated and exposed to spores in 5 mL of saline medium for two days, the volume was then increased to 20 mL and the infection allowed to incubate for 12 extra days. *A. franciscana* were separated into pairs and exposed to spores following the same schedule as *A. parthenogenetica*. The experimental conditions were as follows: the saline medium was sterilized brine diluted to 90 ppt with deionized water, the temperature was 23°C, and feeding was *ad libitum*.

For *E. artemiae*, we also used 22 *Artemia* sampled from the field on 12/3/2015. These individuals were used directly for step (2).

2) We selected spores produced by singly-infected *Artemia* only. After step (1), we ground up each individual *A. parthenogenetica* and each pair of *A. franciscana* in 100 µl deionized water. One quarter of each homogenate was tested for the presence of *A. rigaudi* and *E. artemiae* by PCR (following Rode et al. 2013a). Uninfected or coinfected samples were discarded, while the remainders of the singly-infected homogenates were combined into one *A. rigaudi* sample and one *E. artemiae* sample. These samples were purified on a Ludox density gradient to remove debris as follows. Samples were divided into 2 mL batches, which were layered onto 1.5 mL of a 30 % Ludox (Sigma-Aldrich, USA) solution in 15 mL falcon tubes and centrifuged for 20 min at 7 000 g. We then replaced 3 mL of the supernate with fresh deionized water, vortexed the sample, and re-centrifuged for 4 min at 5 000 g. Finally, we removed 3 mL of the supernate, resulting in a purified and rinsed spore sample.

3) We used the recombined and purified homogenates described in step (2) to infect new populations of lab-bred and parasite-free *Artemia*; these new stock populations thus contained only *A. rigaudi* or only *E. artemiae*. For each microsporidian, the new, 'purified', stock population consisted of ~150 individuals, of which half were *A. parthenogenetica* and half were *A. franciscana*. These hosts were exposed to the spores together in 150 mL of sterilized brine for two days, then the hosts were transferred to large separating funnels and volume was increased to ~2.5 L. The purified stock populations were maintained in sterilized brine diluted to 90 ppt with deionized water, at 23°C and with *ad libitum* feeding.

Whereas the experiment described by Lievens et al. (2018) used spores from both the field-based and the purified stock populations, this experiment only used spores produced by the latter. The *A. rigaudi* and *E. artemiae* inocula used to start the serial passages were obtained by collecting the feces produced by the stock hosts over a 20-hour period. We collected the fecal solutions, homogenized them, and estimated their spore concentration as described by Lievens et al. (2018).

#### Experimental evolution

We serially passaged the microsporidians *A. rigaudi* and *E. artemiae* on the host species *A. franciscana*, *A. parthenogenetica*, or an alternation of the two. After 10 passages, we assayed the infectivity, virulence, and spore production of each line, and compared these among treatments.

*Experimental conditions*

We used lab-raised, parasite-free *Artemia* as experimental hosts for both the serial passages and the final assays. The experiment required a constant supply of fresh hosts; to meet this demand we collected live larvae from stock populations of Aigues-Mortes *A. franciscana* and *A. parthenogenetica* several times a week. Larvae were then raised separately until they could be used for the experiment. We controlled for host age and size as much as possible, by using 6 ± 1-week old, (sub)adult *Artemia* for all passages and for the final assays. The only exceptions were passages P4, P5, and to some extent P8 (the latter for *A. franciscana* only), when episodes of lab-wide mortality depleted stock populations. In these cases, we used hosts of all ages. *A. franciscana* experimental hosts were always a random mix of males and females.

All *Artemia* were maintained at 23°C, in a parasite-free saline medium obtained by diluting autoclaved concentrated brine (Camargue Pêche, France) to 90 ppt with deionized water. *Artemia* were fed a solution of freeze-dried microalgae (*Tetraselmis chuii*, Fitoplancton marino, Spain; concentration: $6.8*10^9$ cells/L deionized water). Stock hosts and hosts being raised for use in the experiment were fed *ad libitum*; assay hosts were fed 0.5 mL three times a week (~0.64 mL corresponds to the maximal ingestible amount for an adult Artemia in two days, Reeve 1963). Host groups in the serial passage experiment were fed at the same rate (20 mL per group per two days). For practical reasons, we did not adjust this quantity to host survival, so that hosts in high-mortality groups were probably slightly better fed than those in low-mortality groups. The serial passages and final assays took place under constant light.

*Serial passages*

We subjected *A. rigaudi* and *E. artemiae* to serial passaging under three evolutionary treatments: '*A. f.* host', '*A. p.* host', and 'Alternating hosts'. In the first two regimes, the parasites encountered only *A. franciscana* or only *A. parthenogenetica*; in the third regime, the parasites encountered alternating passages of *A. franciscana* and *A. parthenogenetica*. Each microsporidian × treatment combination was replicated four times, producing a total of 24 parasite lines.

Parasites underwent ten serial passages, each lasting three weeks. To inoculate the first passage, P1, we used spore doses which were calculated to produce a maximal level of infection (as measured by Lievens et al. 2018): 3 000 spores/individual for *A. rigaudi* on both hosts and for *E. artemiae* on *A. franciscana*; 8 000 spores/individual for *E. artemiae* on *A. parthenogenetica* (10 000 spores/individual would have been more appropriate, but *E. artemiae* spores were limiting). For each parasite line, a group of 40 hosts was first exposed to spores in a limited volume of saline medium (300 mL). After two days, the volume was increased to 750 mL, and the infection was allowed to incubate for the rest of the passage time (19 days). After this incubation period, the spores produced by the surviving P1 hosts were used to infect the second passage, P2, as follows: for each parasite line, adult P1 hosts were placed in a strainer which allowed feces (and spores) to escape, but not the hosts themselves. This strainer was suspended over the 40 new (P2) hosts in 1000 mL medium for 2 days. The strainer was then removed, the P1 hosts counted and stored in 96 % ethanol, and the P2 hosts placed in 750 mL fresh medium to incubate the infection for the rest of the passage time (19 days). Infections from P1 to P2 thus occurred directly from old to new hosts (a similar method was used by Rode et al. 2013a, and Lievens et al. 2019). This method was repeated for P3 to P10. On occasion, a high proportion of the new hosts died during the two days of exposure to the old hosts. To prevent stochastic loss of parasite lines, if 5 or fewer new hosts survived, we added five additional new hosts to the group at the beginning of the incubation period (this slightly over-estimated host survival). If all new hosts died, the transmission step was repeated for that line.

Two aspects of our passaging protocol should be pointed out: first, the time between passages (three weeks) is enough to allow infections to be transmitted between hosts (Rode et al. 2013a). Second, we did not control the number of spores that were transmitted from one group of hosts to the next. In all passages after P1, the size of the inoculum depended on the parasite load of the old hosts. These two aspects meant that the lines were allowed to develop their own infection dynamics, just as they would in the field. Any stochasticity caused by variation in these demographic processes is explicitly included in the experiment. At three points in the experiment, we estimated the microsporidian population size at the time of transmission by counting the spore load in the surviving hosts. These counts were done for the P1, P4, and P7 hosts. After passaging (to P2, P4 and P8, respectively), half of each line's hosts were rinsed and ground up in 1 mL deionized water. We then added 429 µL of pure Ludox (Sigma-Aldrich, USA) and vortexed well, resulting in a homogeneous 30 % Ludox sample. Each sample was centrifuged for 15 min at 10 000 g, after which 1 179 µL of the supernate was removed. To rinse the sample, we added 750 µL deionized water, centrifuged for 4 min at 7 000 g, and removed 900 µL of the supernate. The remaining 100 µL contained the purified spores, which we stained with 1 µL 1X Calcofluor White Stain (18909 Sigma-Aldrich, USA); we then counted the number of spores in 0.1 µL using a fluorescence microscope (see Lievens et al. 2018 for details). We estimated the microsporidian population size as the total number of spores in the body of all hosts = the spore count * 2 (because we only used half of the surviving hosts) * 1 000 (because we counted only 0.1 µL of the sample); this measure should reflect the size of the infective population (Refardt and Ebert 2006).

After the transmission step between the P6 and P7 hosts, we collected a backup spore sample for each parasite line. We kept the surviving P6 hosts from each line in 150 mL containing 10 mL algal solution for one week, after which we collected their feces and stored them at 4°C. Because storage at 4°C likely induces bottlenecks, this cannot be seen as an entirely representative snapshot of the microsporidian population at P6, but the alternative ('storage' in live hosts) would have caused further evolution in the samples. For the line *E. artemiae* × '*A. f.* host' – Replicate 4, feces were collected from the P5 hosts instead of the P6 hosts, because the P6 passage had to be skipped (see Results).

##### *Final assays*

At the end of the serial passage experiment, we tested the infectivity, virulence, and spore production of each evolved line in both *A. franciscana* and *A. parthenogenetica*. We tested all surviving parasite lines based on the spores they produced at the end of P10. We also tested a subset of the parasite lines based on backup spores collected after P6, which we call the 'revived' lines (see Results and Table 1).

To collect spores from the P10 hosts (after they had incubated the infection for three weeks), we placed each evolved line's hosts in a conical container containing 80 mL saline medium and 0.32 mL algal solution per individual (the maximum amount one adult Artemia can ingest over a 20 hour period, Reeve 1963), and allowed their feces to accumulate at the bottom of the container. After 20 hours, we removed the hosts and collected 4 mL of fecal solution. The fecal solution was homogenized, and a 100 µL subsample was used to estimate the spore concentration (after dilution to 1 mL with 900 µL deionized water). Homogenization and spore counting were carried out as described by Lievens et al. (2018).

Spores collected from the P6 hosts were not used directly in the final assays because spore survival at 4°C is not indefinite (Lievens et al. unpublished data). Instead, the spores were used to infect a set of 'revival' hosts 6 weeks after their collection, allowed to incubate for three weeks, and re-stored at 4°C. First, we quantified the spore concentration in each P6 sample by homogenizing, staining, and counting a 1 mL subsample as previously described for single samples (Lievens et al. 2018). We then infected a group of 20 hosts (*A. franciscana* for all revived lines except *E. artemiae* × '*A. p.* host' – Replicate 3) with 60 000 spores and allowed the infection to incubate for three weeks. Revival hosts were 5 ± 1 weeks old, and were kept in 250 mL of saline medium under the same conditions as experimental hosts. After three weeks, the revival hosts were placed in conical containers containing 100 mL saline medium, and feces were allowed to accumulate for one week. Accumulated feces were collected and stored at 4°C until the time of the final assays (10 days). At this time, we concentrated the fecal solutions to 4 mL by centrifuging them at 5 000 g for 8 min and removing the supernate; the 4 mL fecal solutions were then homogenized and the concentration measured as above.

A first assay tested the infectivity of each evolved line. For each line, we exposed 40 uninfected *A. parthenogenetica* and 40 uninfected *A. franciscana* to a low dose of the spores produced by the P10 or revival hosts. The assay hosts were placed in individual hemolymph tubes containing 1.5 mL saline solution and 750 spores; this dose was designed to produce a general level of 20-25% infection while being comparable across treatments (Lievens et al. 2018). After two days in this low volume, we could be sure that hosts had ingested all of the spores (Reeve 1963), and a further 2.5 mL saline medium was added to the tube. The infection was allowed to incubate for five additional days, after which surviving individuals were sacrificed and tested for infection by PCR (Rode et al. 2013a). Seven days should not be enough time for differential mortality to affect the outcome of infectivity (Lievens et al. 2018).

In parallel with the first, a second assay tested the virulence and spore production of each evolved line. For each line, we exposed 40 uninfected *A. parthenogenetica* and 40 uninfected *A. franciscana* to a saturating spore dose, and tracked the mortality and spore production of these individuals for 90 days. The assay hosts were exposed in individual *Drosophila* tubes containing 1.5 mL saline solution and 3 000 spores; this dose was designed to produce a maximal level of infection while remaining comparable across treatments (Lievens et al. 2018). Because we suspected that this dose would lead to fewer infections when *A. parthenogenetica* was exposed to *E. artemiae*, we increased the level of replication for these combinations, exposing an additional 20 uninfected *A. parthenogenetica* to the parasite. We also included 80 control *A. parthenogenetica* and 80 control *A. franciscana*, which were not exposed to spores but otherwise treated identically. After two days, 10 mL saline solution was added to the tube, and the infection was allowed to incubate until the host's death or until 90 days had passed. The saline medium was replaced once a week. Deaths were noted daily, and dead individuals were collected and stored in 96 % ethanol. Surviving individuals at the end of the assay were sacrificed and stored in 96 % ethanol. All individuals were then tested for infection by PCR following Rode et al. (2013a). Since the PCR tests were sensitive to false negatives (DNA of dead individuals could degrade before collection, and parasite load could be strongly reduced by the end of the assay), we also looked for spores in a fecal sample from each 'negative' individual. The fecal samples had been collected on days 10, 16, or 37 and stored at 4°C, as described in the next paragraph. As a consequence, we could be sure of the infection status of all individuals that died on or after day 11 (i.e. after the first spore collection period). Before that date, we could not exclude false negatives.

In the second assay, spore production was measured by quantifying the number of spores produced by infected individuals over a two-day period after three weeks of infection; this corresponds to the window for transmission during the serial passage experiment. Two days after changing the saline medium on day 21 (i.e. on day 23), we collected 800 µL of each host's feces (containing spores) from the bottom of their tube. These samples were stored at 4°C until the infection status of each individual was known. Then, for each line, we pooled all fecal samples produced by infected *A. franciscana* and *A. parthenogenetica*. We quantified the spore count of 1 mL subsamples as described in Lievens et al. (2018). By pooling samples, we minimized the error introduced by the sampling method, which is probably quite variable (since it depends on e.g. the consistency of the feces, the rate at which the host disturbs them, etc.). Each sample was counted twice; the two counts were averaged during the analyses.
